# ADTSIM 2.0: An updated simulation of American dog tick (*Dermacentor variabilis* Acari: Ixodidae) population dynamics

**DOI:** 10.64898/2026.11.20.768101

**Authors:** Shania Sanderson, MyKenna Zettle, Oleksii Dubovyk, Chi Wei, Alexandra Cumbie, Joshua B. Benoit, Sadie J. Ryan, Angela James, Holly Gaff

## Abstract

The American dog tick, *Dermacentor variabilis* (Say), is found across most of the central and eastern portions of North America, and is a known vector of human and veterinary pathogens. While this tick has been found throughout this region for decades, there is still little understanding of the drivers for the highly variable annual densities. To explore the relative effects of hosts, environmental conditions, and tick biology, we introduce ADTSIM 2.0, an upgraded version of the ADTSIM model (Mount and Haile 1989) that simulates the lifecycle of *D. variabilis*. The model uses a weekly time step and follows the life stages (egg, larvae, nymph, and adult), questing behavior, host-seeking dynamics, and survival rates under varying environmental conditions. The model includes calculations for tick interactions with small- and medium-sized mammal hosts and factors in critical environmental influences such as temperature on tick development and questing activity. It addresses the effect of questing behavior on tick longevity, emphasizing the trade off between survival and host-seeking. The results show that accurate estimates of both host densities and weather information are critical to predict local *D. variabilis* densities. This model provides the first step to building a tool for public health information, offering a framework for assessing tick control measures and is adaptable to add tick-borne pathogen dynamics for both human and animal diseases.

## Introduction

Ticks are obligate parasites that are capable of transmitting a greater variety of pathogens compared with any other arthropod vector (Pfäffle et al. 2013). Tick-borne diseases directly effect morbidity and mortality of humans and animals worldwide, and are a growing concern due to a number of factors including climate change, habitat destruction, and altered tick-host-pathogen dynamics (Beard et al. 2021). Many tick species, such as the American dog tick, *Dermacentor variabilis* (Say), are predicted to undergo a significant range expansion in the future (Minigan et al. 2018). *Dermacentor variabilis* is a medically-relevant tick species found from central to eastern North America (Lado et al. 2021), and is a vector of several pathogens including *Rickettsia rickettsii*, the causative agent of Rocky Mountain spotted fever (RMSF) (Burgdorfer 1975) in humans, and *Anaplasma marginale* Theiler, the causative agent of bovine anaplasmosis (de la Fuente et al. 2001) in cattle.

More than 1,000 cases of RMSF and other spotted-fever group rickettsioses are reported in the U.S. every year, highlighting a growing need for increased control, surveillance of tick-borne diseases, and accurate modeling of tick distributions (CDC 2024). While only a small number of cases are reported annually for bovine anaplasmosis (Villar et al. 2023), the economic consequences of an outbreak can be significant for the cattle industry. Preventing tick bites and reducing the spread of tick-borne diseases has been an ongoing challenge. However, methods for tick control commonly target other species of veterinary or agricultural importance, and wide-scale control of *D. variabilis* has been poorly studied (White and Gaff 2018).

Immature *D. variabilis* ticks primarily feed on small mammals, many of which have been found to be competent reservoirs for *R. rickettsii*, including hispid cotton rats (*Sigmodon hispidus* Say and Ord), meadow voles (*Microtus pennsylvanicus* Ord), and white-footed mice (*Peromyscus leucopus* Rafinesque) (Gage et al. 1990; Magnarelli et al. 1983). Medium-sized and larger mammals are targeted by questing adults, and some, such as raccoons (*Procyon lotor* Linneaus) and Virginia opossums (*Didelphis virginiana* Kerr), act as reservoirs for *R. rickettsii* (Zimmerman et al. 1988). Ticks become infected by feeding on infected hosts or through transovarial transmission (Hayes et al. 1980; McDade and Newhouse 1986). After infection, ticks can transmit specific pathogens to uninfected mammalian hosts during feeding or to other ticks by cofeeding (Hayes and Burgdorfer 1989). While much of the *R. rickettsii* cycle is based in wildlife, the majority of *A. marginale* infections are found in domestic cattle with indications that ungulates that may serve as a wildlife reservoirs (Kocan et al. 2010). Since immature and adult ticks have such strong preferences for small and medium (or larger) hosts, respectively, populations of both sized hosts are necessary for sustaining tick populations and pathogens in a specific location (Bishopp and Smith 1938; Smith et al. 1946).

Simulation models offer a cost-effective means to evaluate changes in tick populations and infection rates following different intervention methods, making them valuable for prioritizing prevention strategies that show more potential for the reduction of tick populations. The life history model presented here is a model for *D. variabilis* that modifies and enhances the original ADTSIM model (Cooksey et al. 1990), which was developed to simulate the life history of *D. variabilis* and the transmission dynamics of *R. rickettsii*. The original ADTSIM model took into account a number of biotic and abiotic variables, including host density, fecundity, pathogen transmission rates in hosts and ticks, transovarial transmission, and weather conditions (Cooksey et al. 1990). Our revised version, ADTSIM 2.0, focuses on updates to the tick population dynamics including including finite life stages and limiting the number of ticks per host. There are also updates in several aspects of the model to reflect new information for environmental parameters, phenology, and host dynamics. This model provides the platform for future models to add pathogen dynamics, but here we focus on the complex relationship between the tick biology, host dynamics, and environmental conditions. An understanding of the future risk of tick-borne diseases and potential control methods begins with an investigation of the effects of both biotic and abiotic drivers on tick populations and pathogen transmission.

## Methods

### Overview of changes to original ADTSIM model

The population dynamics model is heavily based on the mechanisms outlined in Mount and Haile’s American dog tick population model (Mount and Haile 1989) and was recoded using R statistical software (R Core Team 2026). The tick population consisted of weekly cohorts of individuals, each belonging to one of the following stages: eggs, host-seeking larvae, on-host larvae, engorged larvae, host-seeking nymphs, on-host nymphs, engorged nymphs, host-seeking adults, on-host adults, and engorged females. The biggest change from the original model is the addition of a activity-dependent maximum lifespan for host-seeking life stages and the addition of a maximum number of ticks per host. Because there has been a paucity of research on *D. variabilis* in recent decades, the majority of parameters in this update are retained from the original model.

### Model structure: life cycle

Similar to the original model (Mount and Haile 1989) and LYMESIM 2.0 (Gaff et al. 2020), ADTSIM 2.0 utilizes a set of age-structured equations with a discrete weekly time step as shown in Figure 1. All age classes contain both male and female ticks in equal proportion except the final engorged adults class, which tracks only females. Following the original ADTSIM, the adults are assumed to have a constant sex ratio of one-to-one with assumed mating on host. Each week of the model simulation, each cohort of each age-class is updated based on survival within that stage or success in advancing to the next stage. This advancement is dependent on temperature for fecundity and development rates of eggs, engorged larvae, engorged nymphs, and engorged females, but engorgement time is kept constant. Survival on-host is density-dependent, and off-host survival depends on habitat, temperature, and saturation deficit. Host-finding rates are calculated based on host density, temperature, and day length.

**Figure 1:**
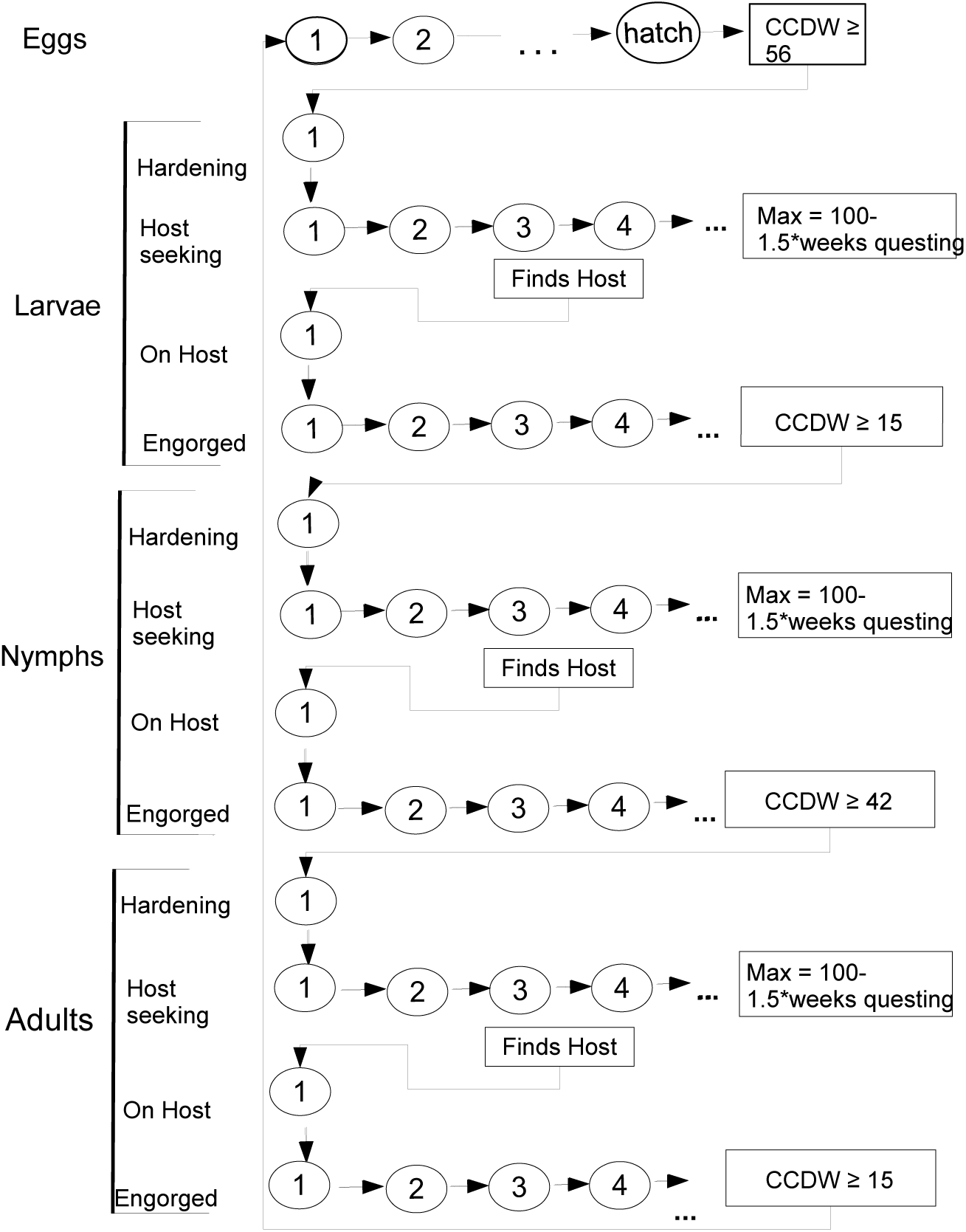
Life cycle model for *Dermacentor variabilis*. Each circle is a weekly age cohort subjected to a mortality rate influenced by parameters such as the stage, activity, weather conditions, and host availability. The Cohort Cumulative Degree Week (CCDW) is a variable assigned to each cohort that tracks the cumulative degrees by which the weekly average temperature exceeds the developmental threshold temperature (DT), set at 10 for eggs and larvae and 9 for nymphs and adults. An egg cohort transitions to larvae when its CCDW exceeds 56, while an engorged larval cohort emerges as nymphs when its CCDW surpasses 15. Similarly, an engorged nymphal cohort will develop into adults once its CCDW is greater than 42, and an engorged adult female cohort will lay eggs when the CCDW exceeds 15. Each host-seeking stage cohort is limited to a maximum of 100 weeks, reduced by 1.5 times the number of weeks the cohort has been questing without successfully finding a host.

Meadow voles and hispid cotton rats are commonly reported hosts of immature stages while opossums, raccoons, and dogs (*Canis lupus familiaris* Linnaeus) are hosts of the adult stage of *D. variabilis* (Gage et al. 1990; Kidd et al. 2006; Magnarelli et al. 1983; Zimmerman et al. 1988). The hosts are modeled as two groups: small-sized mammals (SMM) and medium-sized mammals (MSM), which is identical to the original ADTSIM model. Following the original ADTSIM modeling we assume a constant host density with a ratio of 20:1 for SMM:MSM, and we also use the original constant birth and death rates that yield an average life span of approximately three months for SMM and one year for MSM. The impacts of host life span and ratio of host types will be explored in future models that include pathogen dynamics.

### Model structure: temperature-dependent development and fecundity rates

Similar to the original ADTSIM, the rates of hatching and molting are contingent upon temperature. Cohort-cumulative degree weeks (CCDW) represents the total accumulation of degrees per week above the developmental threshold temperature (DT) for each age-class cohort. If the difference between the weekly mean temperature and the critical temperature for the specific stage is positive, that difference is added to the CCDW for each age-class cohort, but if the difference is zero or negative, the CCDW remains unchanged. The critical temperature for eggs and engorged larvae is +10°C, and +9°C for engorged nymphs and engorged adults.

Using the development times from the original model that came from on the work of Smith et al. (1946). When the CCDW is greater than 56 for any cohort of eggs, the eggs are then transitioned into the hardening larval stage. For engorged larval cohorts, reaching a CCDW of 15 or more advances them to the hardening stage of nymphs. Likewise, reaching a CCDW of 42 or more for engorged nymph cohorts advances them to the hardening stage of adults. Finally, reaching a CCDW of 15 or more for an engorged female cohort produces a cohort of eggs.

We utilize the temperature-dependent quadratic equation for fecundity, as derived from regression analysis of Campbell and Harris data reports (Campbell and Harris 1979) in the original model (Mount and Haile 1989). The number of eggs laid by a female is given by:

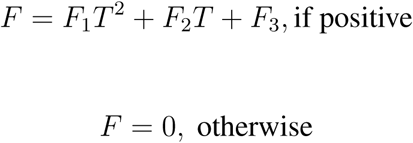

where *T* is temperature and *F*_1_*, F*_2_*, F*_3_ are parameters (See Table S6). This equation gives positive values for the number of eggs laid with a weekly average temperature between 12 and 38°C.

### Model structure: survival rates off-host

Similar to the original model, we employ the habitat classification system proposed by Sonenshine and Levy (1972) to categorize different habitat types (forest, *F*; forest-meadow ecotone, *E*; and meadow, *M*) within our model initialization. The ADTSIM 2.0 model used 90% ecotone, 5% forest and 5% meadow. The maximum survival rate for each habitat type is given in Table S2. Furthermore, the base survival rate within a mixed-habitat area is calculated as a weighted average, which depends on the proportion of the tick population inhabiting each specific habitat (Mount and Haile 1989). The baseline weekly survival was then adjusted as a quadratic function of mean weekly temperature and saturation deficit, and the equation and parameters are given in Table S3. Both baseline survival and coefficients for climatic effects differed for each stage and were taken from the original model (Mount and Haile 1989).

### Model structure: activity-dependent maximum survival

Similar to LYMESIM, a maximum lifespan for each life stage was added to this model for several reasons. First, the original Mount-Haile model suggested age-dependent survival rates with base, i.e., maximum given the climatic conditions, survival rates lower after each step but never reached zero (Mount and Haile 1989). In theory, this results in no maximum age threshold for any individuals, and it was possible, although unlikely, that some individuals would have been able to survive for unrealistically long durations of time under suitable climatic conditions. Second, we introduced a questing penalty for host-seeking individuals. Finding a host is required for survival, but it exposes ticks to dry air outside of the soil and causes them to lose body water and burn through their limited fat reserves (Rosendale et al. 2017), thus decreasing the maximum potential lifespan.

To address this issue, the model has a maximum age limit of 100 weeks for each host-seeking cohort for all life stages, along with a questing penalty of 1.5 weeks for every week spent questing. This approach assumes that ticks possess finite energy reserves, allowing survival for up to 100 weeks without activity, but with active questing consuming 1.5 times the energy expended during periods of inactivity similar to LYMESIM 2.0 (Gaff et al. 2020). The maximum lifespan of 100 weeks was selected based on the evidence for multiple year survival of adults *D. variabilis* under some field conditions Yunik et al. (2015) and longevity here in laboratory conditions (unpublished data).

### Model structure: host finding rates

The host-finding rates (HFRE) for immature ticks (larvae and nymphs) are distinct from those of adults and are accordingly modeled. These rates are contingent upon both temperature (T) and day length (L), with each type of host (SMM or MSM) having a respective host finding rate (SMM BHFR and MSM BHFR) (Table S4) influenced by their respective base host finding rate (BHFR) and host density (SMM HD and MSM HD). The default values for the parameters *P*_1_*, …P*_8_*, Q*_1_*, …, Q*_8_ are given in Table S7.

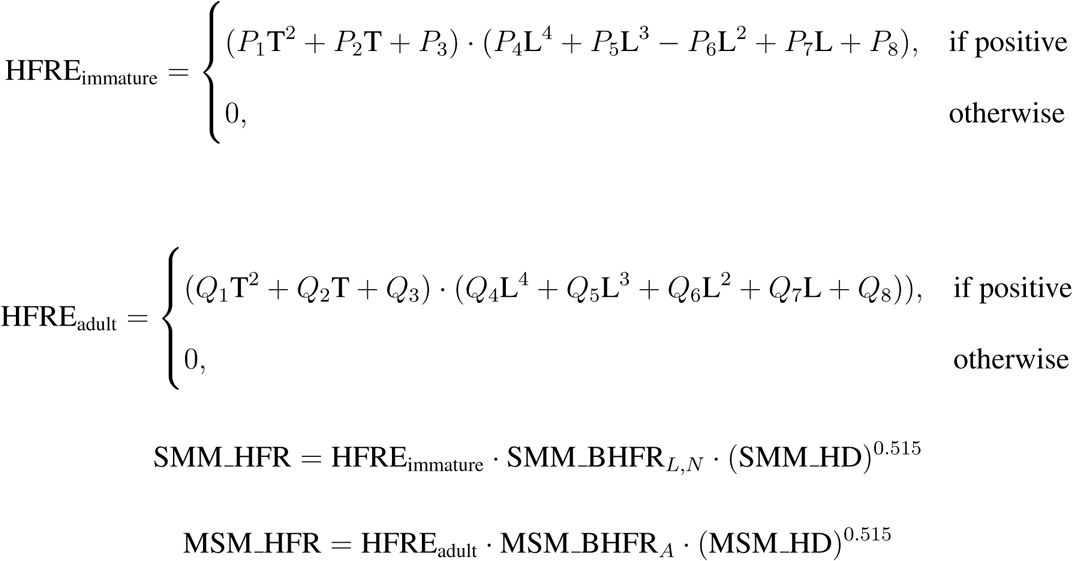

On-host survival is modeled as a relationship with density (ticks/host) as shown in Figure 2 for each life stage. Survival values change linearly with tick density within the threshold range but stay constant for densities outside these limits. This framework was adapted from the original simulation (Mount and Haile 1989). Additionally, there is now a maximum capacity for ticks per host that limits the number of ticks by life stage and host type. These limits are estimated from the values used in LYMESIM Gaff et al. (2020) and field data Cumbie (2020); Rose et al. (2018) (see Table S5).

**Figure 2:**
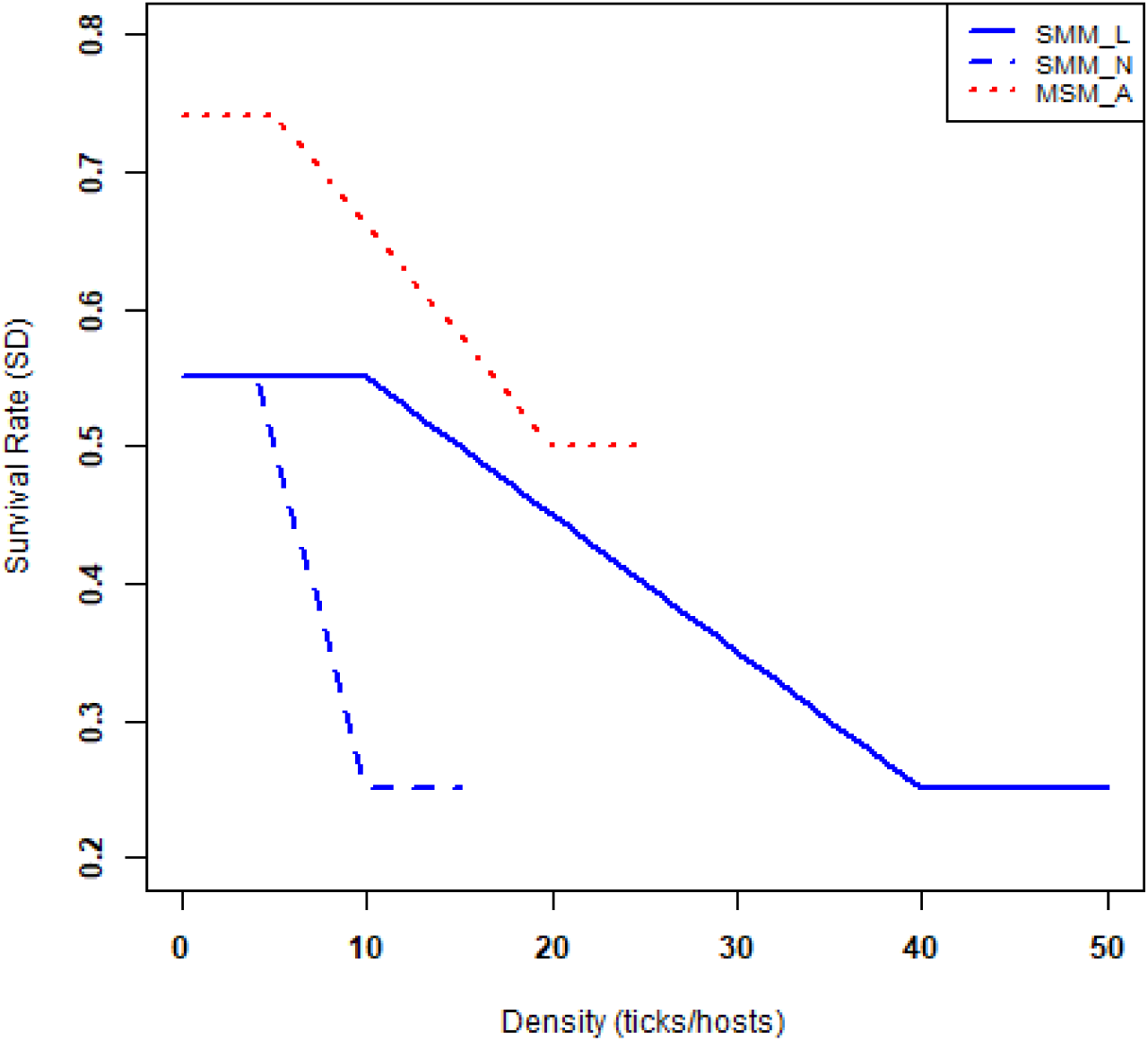
Representation of the relationship between weekly survival of immature (blue) and adult (red) ticks on hosts (SMM and MSM, respectively) versus tick density (ticks/host), adapted from Mount and Haile (1989)

### Output variables

For each model run, three output variables are recorded for each week: surplus adults, larvae on host (LOH), and nymphs on host (NOH). Surplus adults are defined as the total number of adults that are host-seeking in a given week that do not find a host. This surplus represents the adult *D. variabilis* ticks that are likely to be collected during active field surveillance, e.g., flagging. Immature *D. variabilis* ticks are more likely to feed on SMM than humans, thus making them far less likely to be collected via flagging. Instead, immature tick data is measured as the total number of ticks on SMM hosts each week of the model run, and these data can then be compared with ticks that are collected from small mammals in the field.

### Weather data

The weather input files for ADTSIM 2.0 incorporate six variables including weekly average temperature (in degrees Celsius), saturation deficit (in millibars), relative humidity (as a percentage), day length (hours), precipitation (mm), and precipitation index. The precipitation index, *PI*(*t*) is calculated as a weighted average of weekly precipitation, *R*(*t*), and the previous week’s precipitation index, *PI*(*t −* 1). Mathematically, the precipitation index is

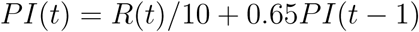

For the initial scenarios of ADTSIM 2.0, the weather input data is the same as used in LYMESIM 2.0 for Norfolk, Virginia, USA (Gaff et al. 2020). This dataset comprised weekly records spanning 2007–2016, supplemented by an average across those years, derived from the North American Land Data Assimilation System (NLDAS) (Cosgrove et al. 2003). Our study explored additional scenario using weather data for 2009-2023 from nearby Chesapeake, Virginia, USA acquired from DAYMET (Thornton and Devarakonda 2011), a 1 km resolution gridded climate product, sourcing multiple input observational products. The “single pixel extraction tool” (Thornton and Devarakonda 2011) was used to obtain and download daily daylength, precipitation, minimum and maximum temperature, and vapor pressure for data spanning between January 1, 2009 and December 31, 2023, for the grid cell that contains the field site. These daily values were used to calculate weekly variables of mean hourly daylength, precipitation (summed), mean daily temperature, and vapor pressure deficit (VPD).

The validation data used for this model is from field data from a long-term field site, CH1, located in Chesapeake, Virginia. This field site is a property managed by The Nature Conservancy adjacent to the Great Dismal Swamp National Wildlife Refuge. It was originally farm fields that were restored starting around 2005, and the property went through succession to return to a closed canopy forest. For additional information on this field site, please see Nadolny and Gaff (2018). Adult *D. variabilis* ticks were collected using a standard flagging technique along a 3000-meter transect monthly from 2010-2023 (ODU Tick Team, unpublished). Additionally, small mammal trapping was conducted at the site from 2010-2018, and immature *D. variabilis* ticks were removed from all trapped small mammals prior to release (Cumbie 2020; Rose et al. 2018). The predicted *D. variabilis* adult surplus using weather data from DAYMET (2010-2016) and NLDAS (2010-2023) was compared with collected flagging data from the field site in the City of Chesapeake, Virginia. The predicted phenological patterns of both immature and adult *D. variabilis* for both weather data sets were also compared the results with field data.

### Scenarios

The model was run using parameter values from the original ADTSIM model. To investigate ADTSIM 2.0, NLDAS average weekly weather data for Norfolk, Virginia, was used to generate outputs of *D. variabilis* immature ticks on host phenology and adult surplus. The results were compared to published (Cumbie 2020) and unpublished data for *D. varabilis* phenology. For year-to-year variation, the simulation was run with similar default parameters, and the weather inputs were from 2007–2016. The scenario for varying hosts was identical to the year-to-year variation, with the addition of varying SMM and MSM populations.

As a further proof-of-concept, we ran an additional set of scenarios that assess how tick populations would be affected by impaired humidity detection (Uhran et al. 2024). This work looked at the function of the Haller’s organ to detect humidity, which is critical to the tick finding suitable locations for survival from desiccation. In summary, this data from this study showed that humidity detection was critical for prolonged tick survival, but it was unknown how this might affect ticks at the population level. Three model scenarios were evaluated by varying the survival rates of questing ticks: (i) a baseline scenario with standard survival rates across all life stages, (ii) a scenario with reduced survival limited to the adult stage, and (iii) a scenario with reduced survival applied to all life stages.

### Sensitivity analysis

For the sensitivity analysis, we assessed the effect of the questing penalty, maximum life span for each host-seeking cohort, base survival rate, and host densities for all host types on adult surplus. To investigate how changes in these parameters affect the results, Latin hypercube sampling (LHS) design was used to conduct a full sensitivity analysis on all parameters. LHS allowed for all parameters to be varied simultaneously (Alden et al. 2013; Marino et al. 2008) at a range of about 10% above and below the initial values found in Tables S1 and S2. The distribution for each parameter range was assumed to be a uniform distribution, and 1000 samples were taken to create the sensitivity parameter sets. The model was then run for each parameter set using average weather data from Norfolk, Virginia, and the results were summarized into three outputs: larvae on hosts, nymphs on hosts, and adult surplus. The partial rank correlation coefficients comparing each parameter to the variation in the output variables were then calculated, and those with a p-value less than 0.05 were considered significant.

## Results

Each model run used the same initial conditions of 300,000 eggs in the first egg cohort, and then the model was run for 10 years using the long-term averaged weather conditions. The “mean long-term” weather data were calculated by averaging weekly parameter values from 2007–2016. With all other parameters held constant, the model was then run using constant hosts with average weather, constant hosts with yearly-varying weather, and varying hosts with yearly-varying weather.

### Scenario: constant hosts, constant weather

To assess the timing of the model predictions for the various life stages, the initial model runs used constant hosts and constant weather (Figure 3). The model was run, and the results converged to a constant predicted tick population as measured by larvae on hosts, nymphs on host, and surplus host-seeking adults for Norfolk, Virginia (Figure 3a and 3c). The model predicted that the peak total larvae on host occurs in the spring, with a limitation at 8000 ticks per hectare due to a maximum carrying capacity for total ticks on hosts, followed by a second peak occurring in the early fall. Similarly, total nymphs on host follow the larval trend at the expected order of magnitude smaller numbers. The total surplus adult populations was predicted by the model to only peak once in the summer. These results mirrored biological expectations as shown in Figure 3b and Figure 3d.

**Figure 3:**
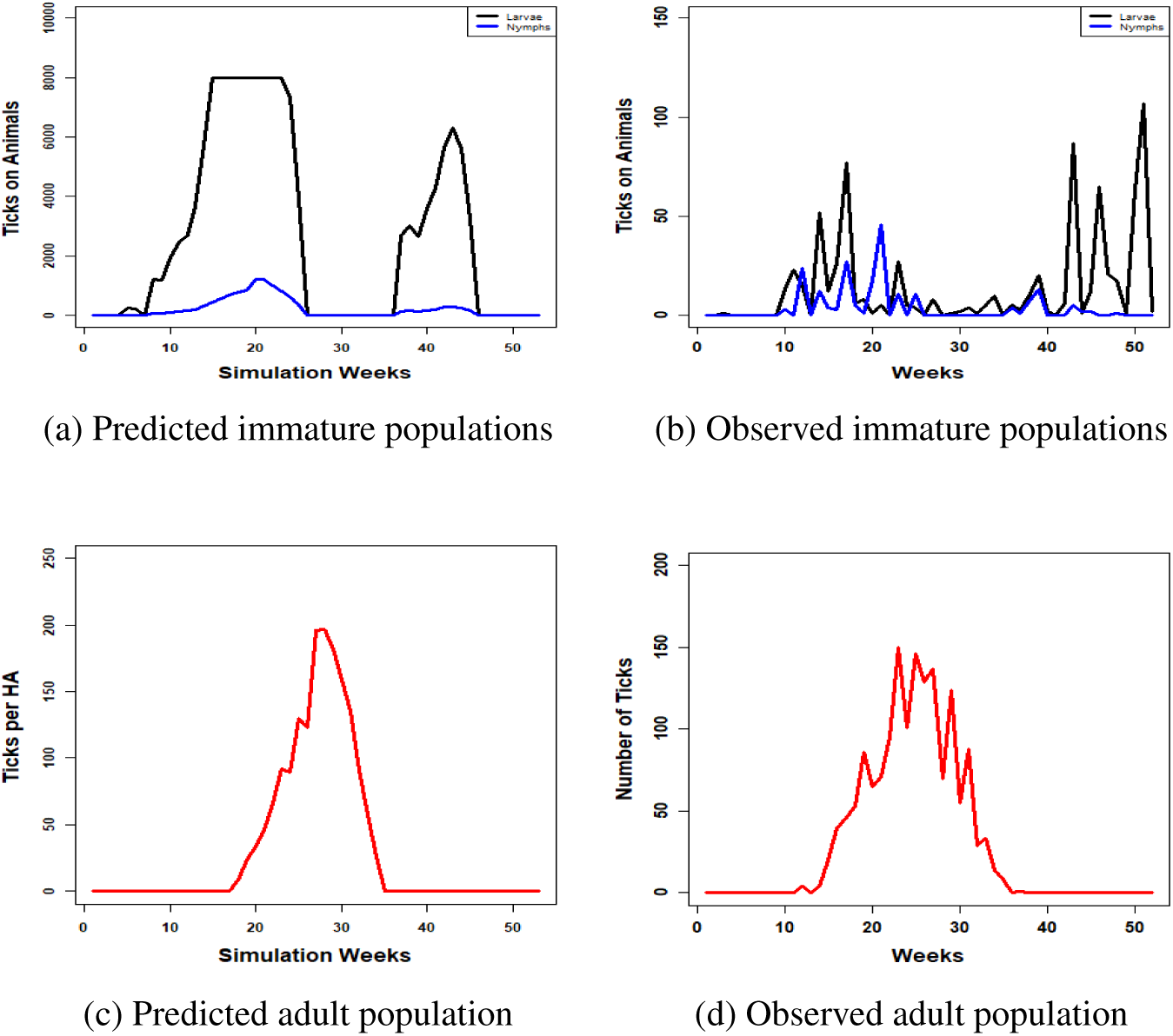
Annual average *D. variabilis* phenology for Norfolk, Virginia. The graphs depict the ADTSIM 2.0 results for (a) immature on hosts and (c) questing surplus adult ticks based on mean weather conditions from 2007 to 2016 in Norfolk, Virginia. These results closely match the overall timing found for field observations of (b) the number of immature *D. variabilis* found on small mammals and (d) the density of adult *D. variabilis* collected via flagging.

### Scenario: constant hosts, variation in weather

The second model run used the year-to-year variable weather data after the initialization period, but the hosts levels were still kept constant. The model results are shown in Figure 4 and show similar trends for the immature life stages, with both larvae and nymphs having spring and fall peaks with the larval spring peak limited by on-host carrying capacity. Surplus host-seeking adult populations fluctuated every summer with a peak in 2010 and a drop in 2013.

**Figure 4:**
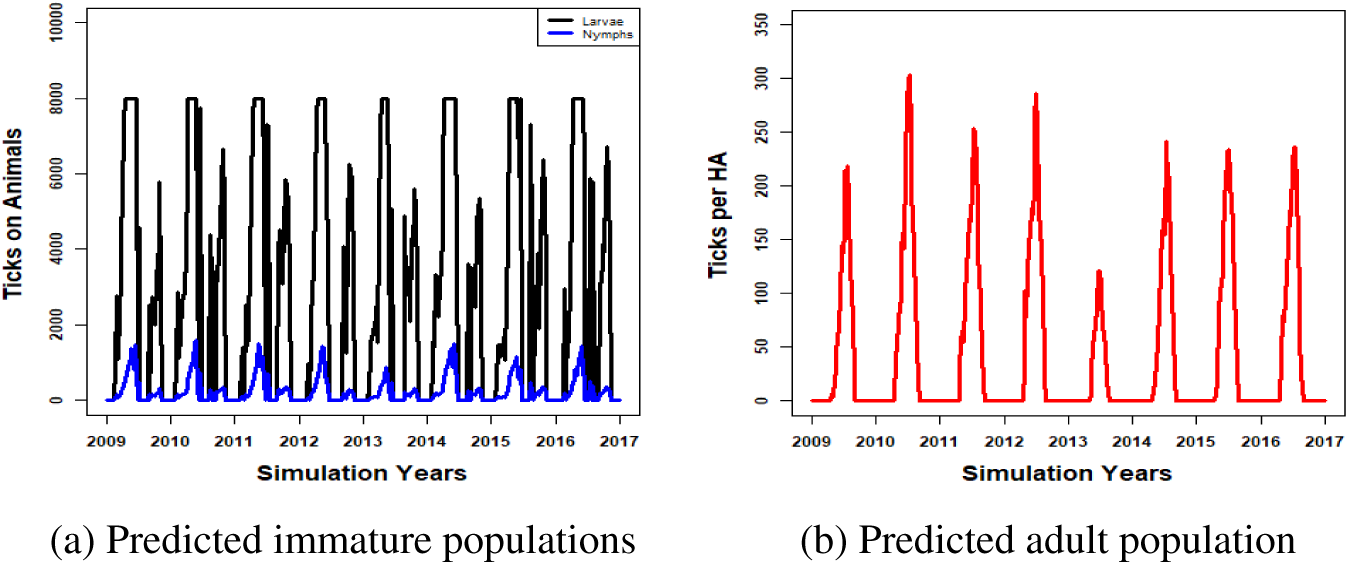
The model results illustrate year-to-year variations in the total number of (a) immature ticks on animals and (b) adult surplus ticks per hectare (HA) based on weather data from 2007 to 2016 for Norfolk, Virginia. Surplus populations of larvae and nymphs exhibit a spike in fall to early spring, while adult populations peak in the summer.

### Scenario: variation in hosts, variation in weather

To compare the model results to a long-term field collection site, the model used both varying weather data and varying host populations (Figure 5). The weather was the DAYMET data for the CH1 field site from 2009-2023, and the results for the model with a constant host population shows a similar phenological pattern to the previous scenario. While the immature ticks on hosts was similar between this scenario and the previous (Figure 4a and 5a), there was variation in the predicted density of host-seeking adults with a much lower predicted maximum density with the new weather data at the CH1 site (Figure 4b and 5b). The host data came from the long-term small mammal collections at that site along with the adult tick collections (Figure 5c). The model output for the year-to-year variation on adult surplus with varying host densities compared to adult *D. variabilis* field data showed similar trends (Figure 5d). Adult populations peaked in the summer of 2011 and decreased annually until 2015 with considerable fluctuations between 2016 and 2023.

**Figure 5:**
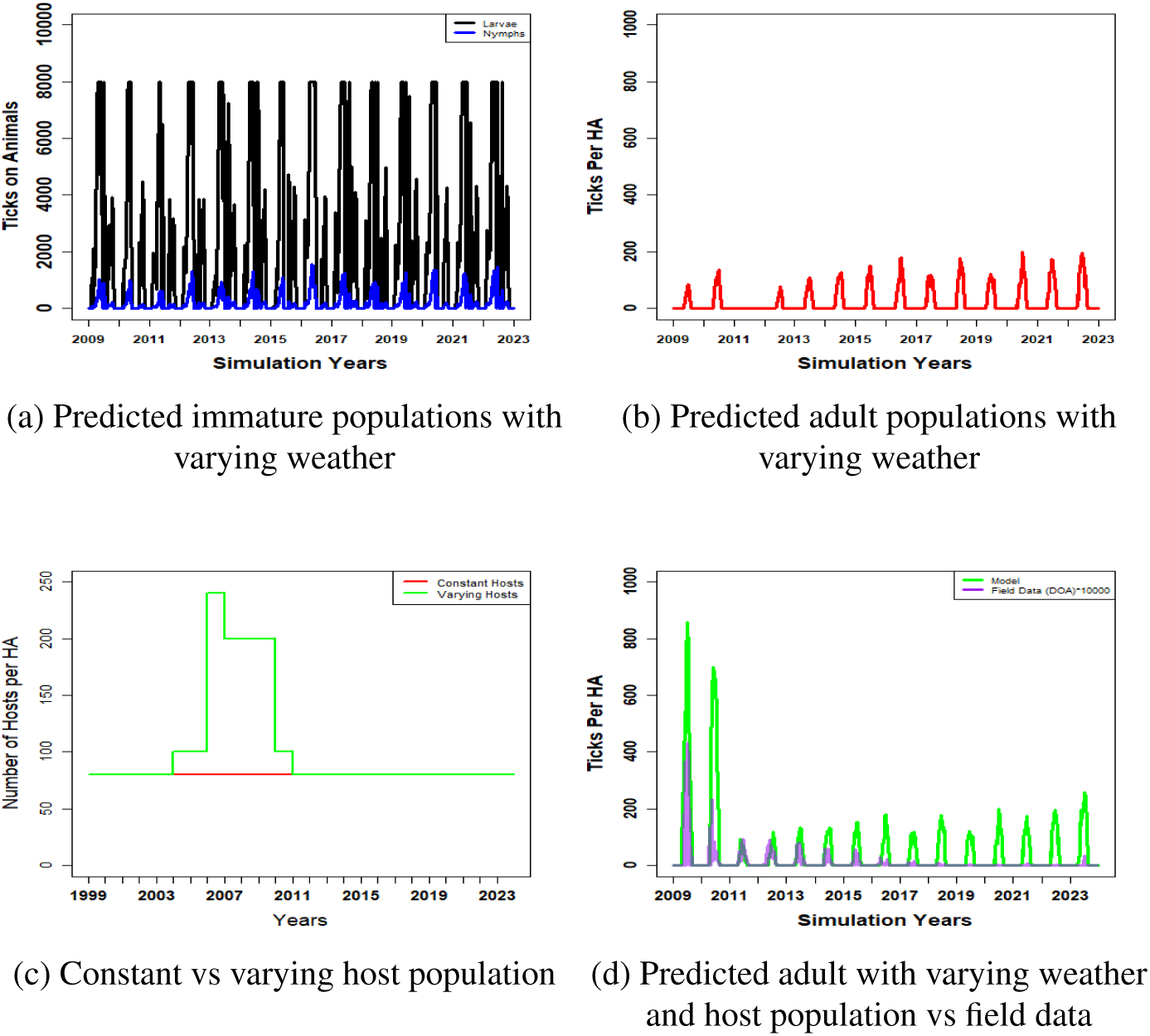
The predicted model scenarios for immature and adult populations (a and b) used varying weather data and constant densities of small-size mammals (SMM) and medium-size mammals (MSM). To account for extreme weather events present in the CH1 data, such as snowstorms and hurricanes, a model scenario for adult populations was run with varying weather and varying hosts (c) and then compared with observed density of adults from long-term field site, CH1 field data (multiplied by 10000) is presented for validation purposes (d). The trends produced by the model closely mirror the observed value trends.

### Scenario: variation in survival

Based on previously reported survival experiments (Uhran et al. 2024), ticks lacking the ability to detect humidity exhibited increased weekly mortality. When this elevated death rate was applied only to adult ticks (assuming similar mortality rates for males and females), the model predicted a significant decline in ticks density (Uhran et al. 2024). Extending the increased mortality to larvae and nymphs did not result in additional population declines beyond those already observed, suggesting that other factors, such as host location, are more critical to their survival. These results suggest that the ADTSIM 2.0 model could be a useful tool to extend specific biological studies and to provide a baseline assessment of whether *D. varabilis* populations may be affected.

### Sensitivity analysis

The results of the sensitivity analyses showed that changes in host parameters and survival rates affected adult populations the most (Figure 6). Sensitivity analyses results revealed that simulation outputs covaried with host finding, fecundity, and egg survival rates as key parameters for immature life stages. Similar parameters were also found to be key for adults with medium-sized mammal base host finding rate for adults being an additional factor (see S8, S9, S10). One of the major changes with this model was the addition of maximum lifespan of 100 weeks and questing penalty, and so the survival of all life stages was calculated. The results showed with the weather conditions in these scenarios that the no ticks were able to survive longer than 60 weeks (Figures S1 and S2).

**Figure 6:**
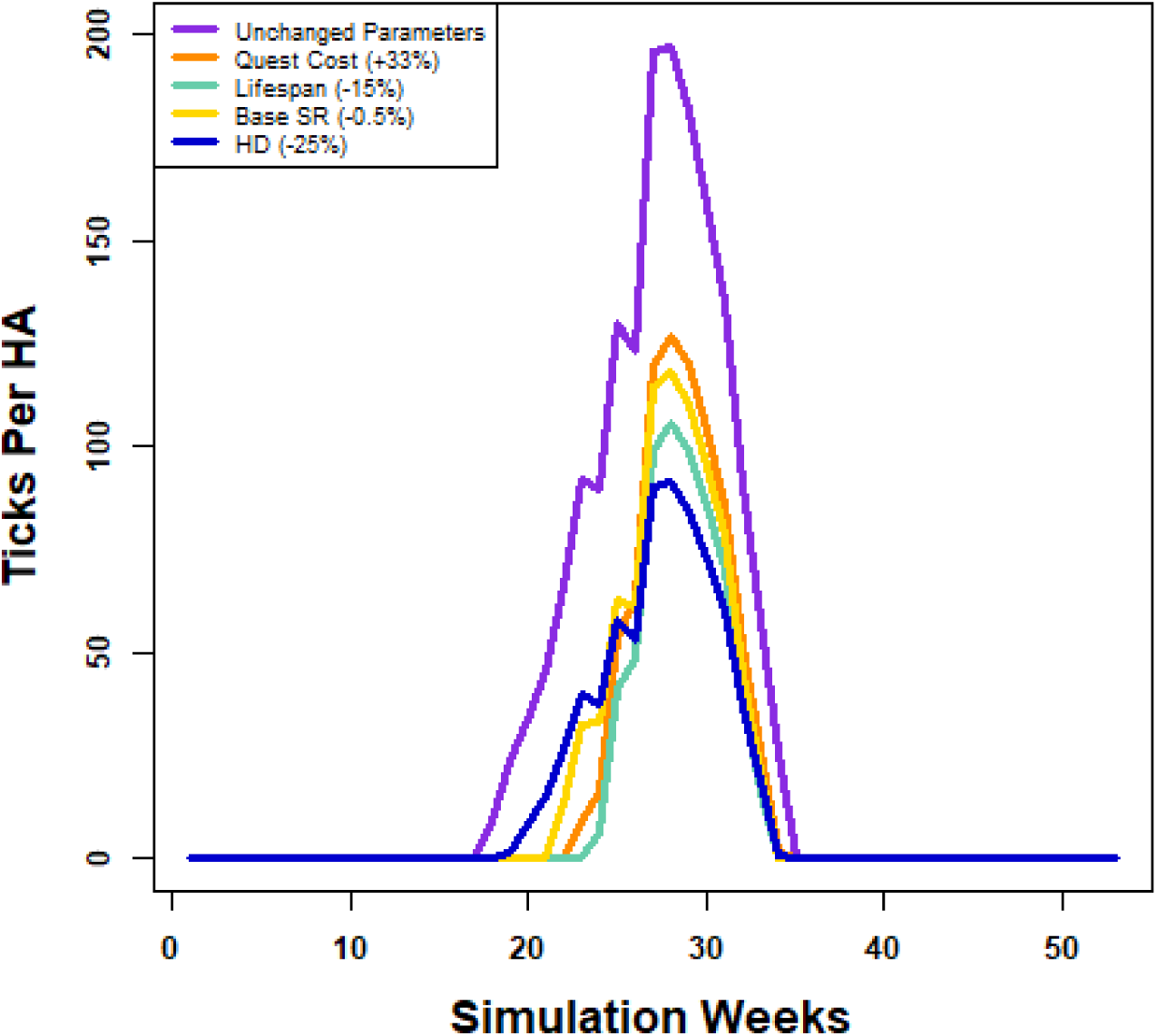
Sensitivity analysis for ADTSIM 2.0 examined the response average yearly surplus density of adult ticks per hectare (HA) to various parameters. The scenarios considered were: (1) the original trends of all surplus ticks without any parameter adjustments, (2) an increase in questing cost by 33% to 2 weeks, (3) a reduction in the overall lifespan by 15% to 85 weeks, (4) a decrease in the basic survival rate (BSR) by 0.5, and (5) a 25% decrease in host density from the original values.

## Discussion

Mathematical models are useful for simulating the complex relationships and interactions presented by ecological systems. Such models can be an effective method for measuring changes in vector populations and infection rates, as well as the effectiveness of different control methods. Building on the model developed by Mount et al. (1989), we have developed ADTSIM 2.0, a model that accounts for the most recent information on *D. variabilis* life history and phenology. Using data published in (Rose et al. 2018), we set the total host-seeking life stage to 100 weeks, allowing populations a longer period to establish, and the questing to 1.5 weeks.

Like all attempts at simulating real-world ecological interactions, ADTSIM 2.0 is not without its limitations. This model was designed to function in a primarily non-anthropogenic system; current host availability, tick-survival, and climate conditions are based on ideal tick habitat. In more urbanized areas, this model would need some considerable adjustments to account for differences in host availability and community dynamics. ADTSIM 2.0 assumes constant host populations, which is an unlikely scenario outside of a simulation. However, the model is designed to easily accommodate variable host dynamics.

The accuracy of ADTSIM 2.0 depends significantly on the fit of the provided climate data, which can vary based on the level of geographical smoothing applied to the data. Furthermore, the size of the location that is being simulated, as well as the scale at which the climate data is collected at, can also result in significantly different model outcomes. We validated our model using data from a field site in Chesapeake, VA, a rural area with observable tick and host populations. However, when we compared the results with data from Norfolk, Virginia, a more urban area located less than 20 miles away, climate data were a much less reliable predictor. The sensitivity analysis results indicate that multiple parameters significantly influence immature and adult populations. This information can aid in developing control methods and has already been utilized to validate experiments related to *D. variabilis* survival. The sensitivity results help to identify the key variables to target with interventions that have greatest potential for reducing tick populations. As an example, this model was used to assess how populations of *D. variabilis* will be affected by increased mortality from impaired humidity detection, which surprisingly showed a substantial effect on adult stages with little reduction in population levels for the juvenile stages (Uhran et al. 2024).

ADTSIM 2.0 serves as a practical tool for further investigation and mitigation of *D. variabilis* populations. Additionally, the development of ADTSIM 2.0 and the final simulation have enhanced the understanding of the nuanced relationships between ticks, their associated pathogens, and preferred hosts. This framework can also be adapted to study other tick species and their respective pathogens.

## Acknowledgments

Funding for this research was partially provided by the United States Department of Agriculture (USDA), Animal and Plant Health Inspection Service (APHIS), Center for Epidemiology and Animal Health award No. AP20VSCEAH00C126. The findings and conclusions presented are those of the authors and should not be taken to represent any official USDA or US government determination or policy. The R code for this version of ADTSIM is freely available via a request directly to the corresponding author.

## Supplemental Materials

**Table S1:** Factors for initialization of ADTSIM 2.0. Values are similar to ADTSIM differing values are indicated with an asterisk(*). Expected host densities vary yearly across ideal *D. variabilis* habitats. Values each group of hosts are reflective from more recent studies.

| **Factor** | **Value(s)** | **Comments** |
|---|---|---|
| Simulation start week | 1 | First week of January of first year |
| Initial tick population | 300,000 eggs | 0 all other age cohorts |
| Habitat Distribution | (0.05, 0.95, 0.05) | Forest, Ecotone, Meadow |
| Small-sized mammal density | 80 per hectare | 0% pathogen prevalence |
| Medium-sized mammal density | 4 per hectare | 0% pathogen prevalence |
| Maximum lifespan per life stage | 100 weeks | Reduced by activity |
| Questing penalty | 1.5 weeks | Life span reduction for activity |
| Length of simulation | Varies | Weekly time step |

**Table S2:**
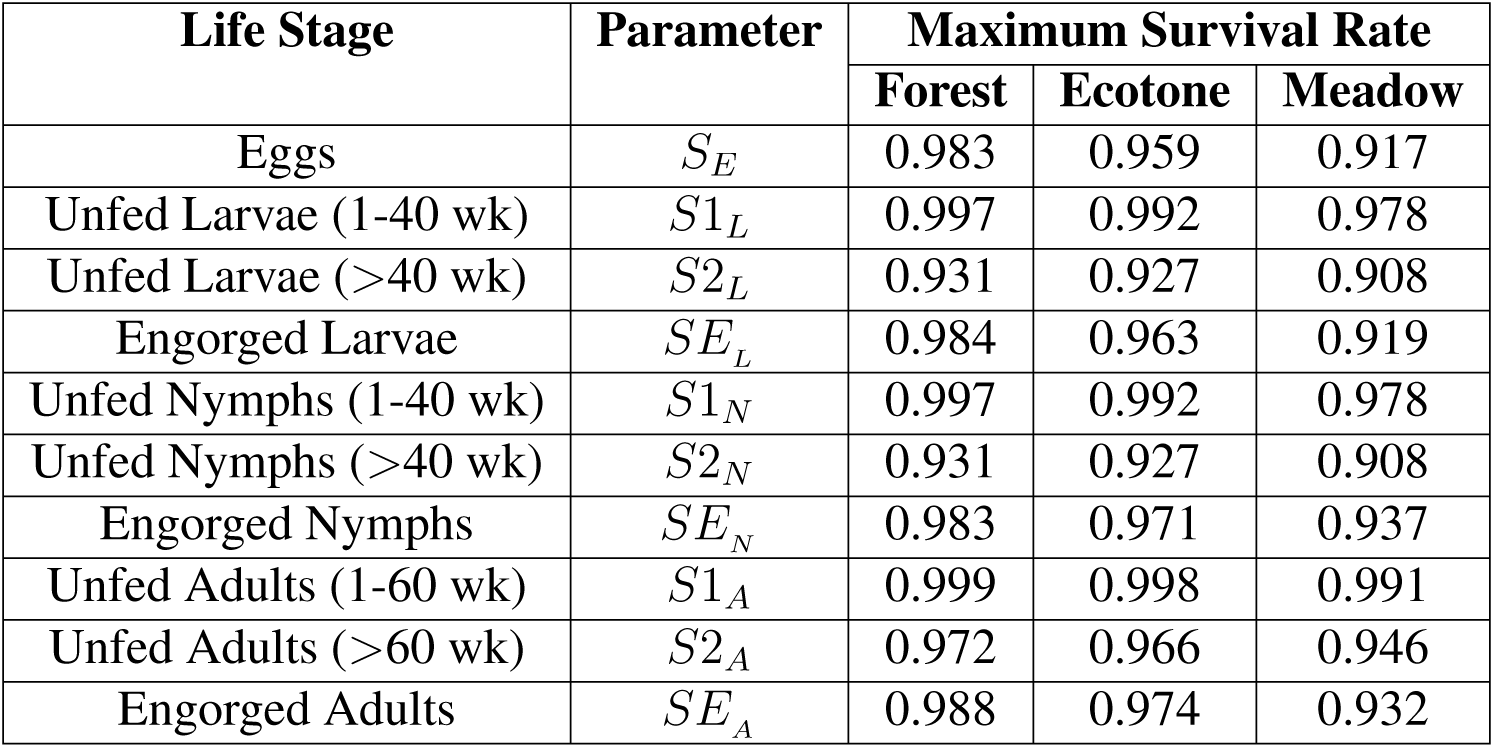
Maximum survival rates for different life stages across habitats. Values are taken from Mount and Haile (1989).

**Figure S1:**
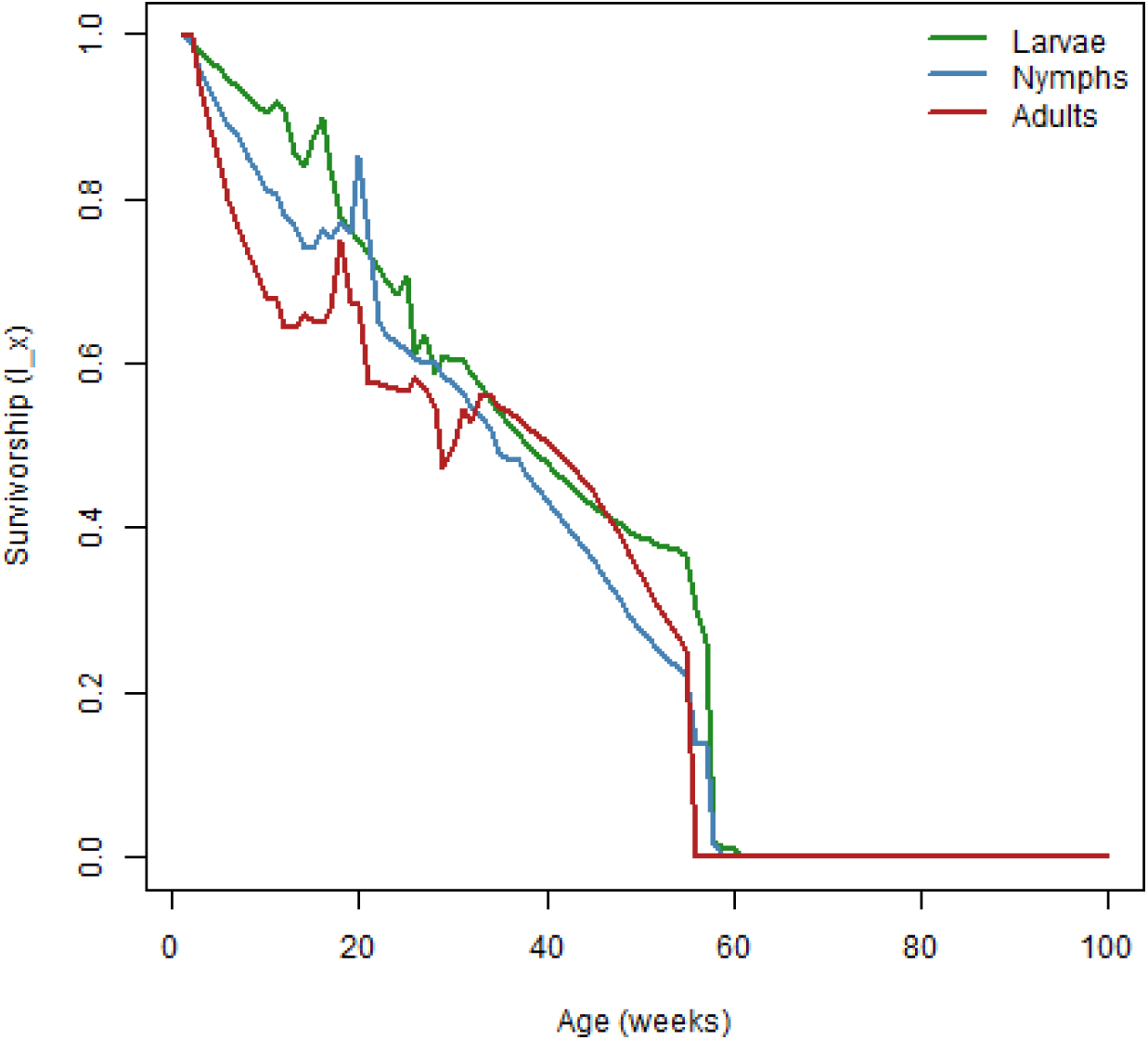
Survivability *Lx* plot for host-seeking immature and adult populations. The model run was with varying weather and constant host densities.

**Figure S2:**
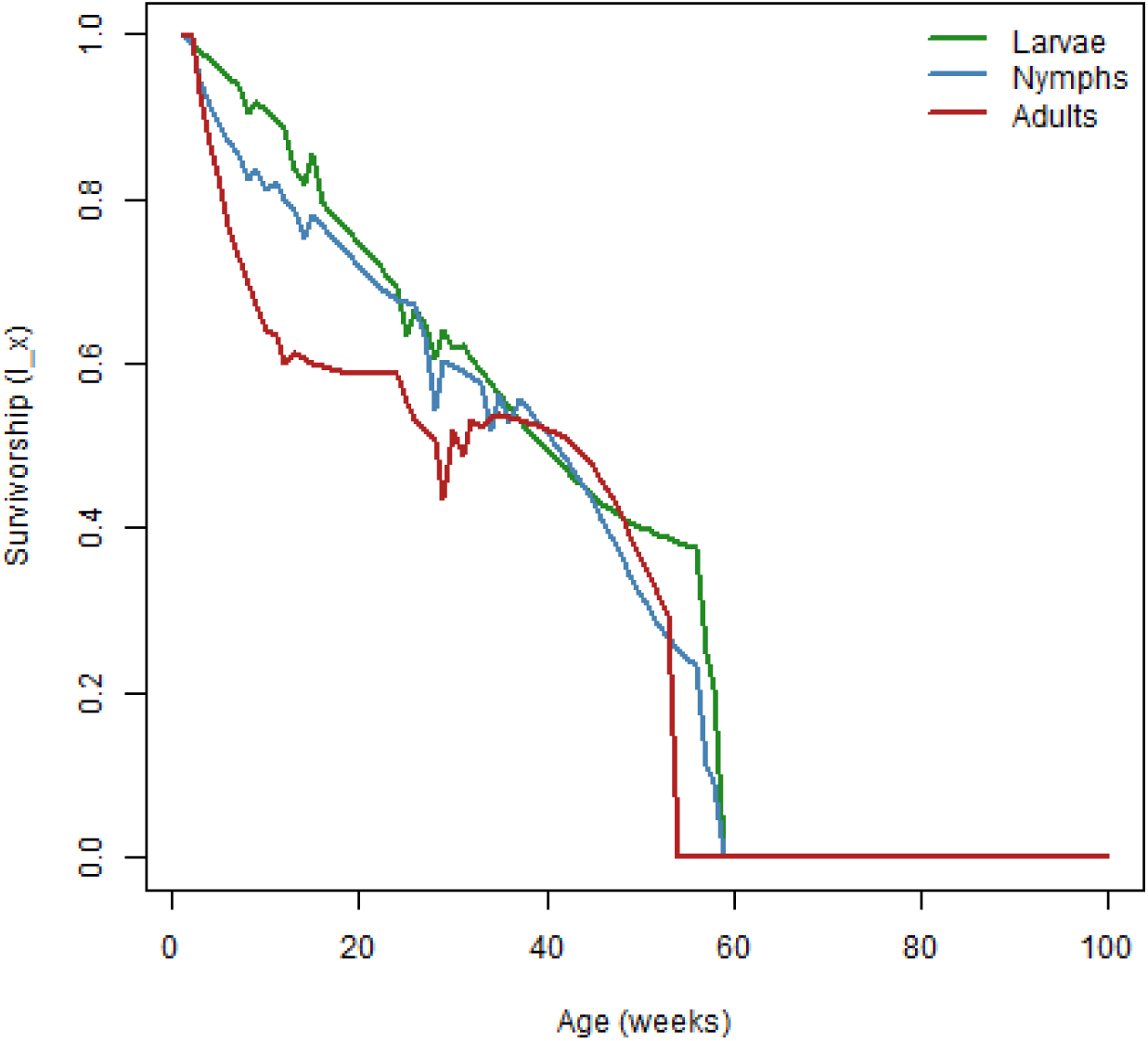
Survivability *Lx* plot for host-seeking immature and adult populations. The model run was with average yearly weather and constant hosts densities.

**Table S3:**
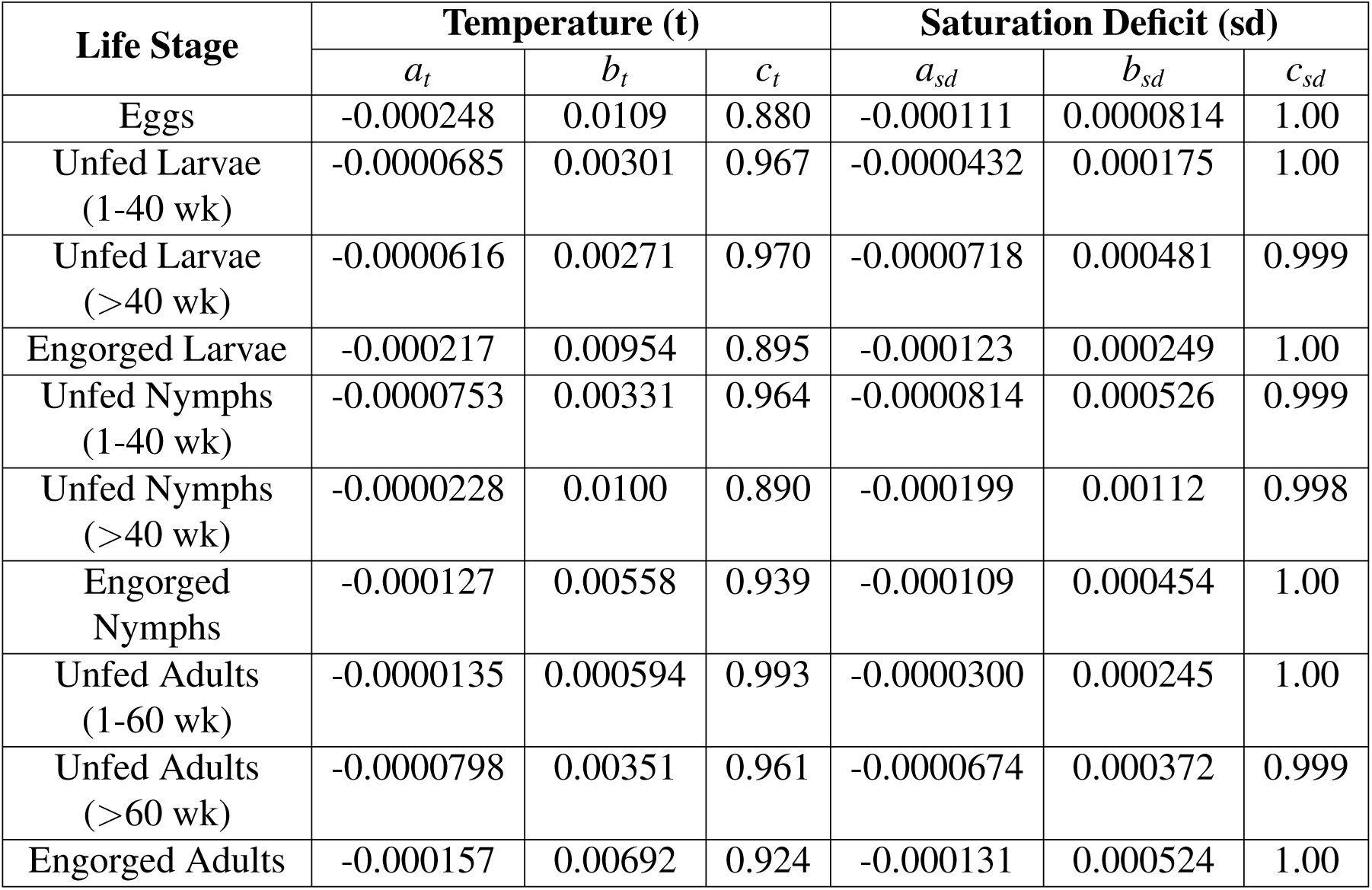
Temperature and saturation deficit parameters are shown for the weather adjustments to survival for different life stages. These parameters are used in the equation to calculate the exact survival for a given temperature (*T*), saturation deficit (*SD*), and maximum survival life stage (*S_x_*) as calculated by the weighted average in Table S2 using: *Surv* = *S_x_*(*a_t_T* ^2^ + *b_t_T* + *c_t_*)(*a_SD_SD*^2^ + *b_SD_SD* + *c_SD_*)

| **Life Stage** | **Temperature (t)** |  |  | **Saturation Deficit (sd)** |  |  |
|---|---|---|---|---|---|---|
|  | *a<sub>t</sub>* | *b<sub>t</sub>* | *c<sub>t</sub>* | *a<sub>sd</sub>* | *b<sub>sd</sub>* | *c<sub>sd</sub>* |
| Eggs | -0.000248 | 0.0109 | 0.880 | -0.000111 | 0.0000814 | 1.00 |
| Unfed Larvae (1-40 wk) | -0.0000685 | 0.00301 | 0.967 | -0.0000432 | 0.000175 | 1.00 |
| Unfed Larvae (>40 wk) | -0.0000616 | 0.00271 | 0.970 | -0.0000718 | 0.000481 | 0.999 |
| Engorged Larvae | -0.000217 | 0.00954 | 0.895 | -0.000123 | 0.000249 | 1.00 |
| Unfed Nymphs (1-40 wk) | -0.0000753 | 0.00331 | 0.964 | -0.0000814 | 0.000526 | 0.999 |
| Unfed Nymphs (>40 wk) | -0.0000228 | 0.0100 | 0.890 | -0.000199 | 0.00112 | 0.998 |
| Engorged Nymphs | -0.000127 | 0.00558 | 0.939 | -0.000109 | 0.000454 | 1.00 |
| Unfed Adults (1-60 wk) | -0.0000135 | 0.000594 | 0.993 | -0.0000300 | 0.000245 | 1.00 |
| Unfed Adults (>60 wk) | -0.0000798 | 0.00351 | 0.961 | -0.0000674 | 0.000372 | 0.999 |
| Engorged Adults | -0.000157 | 0.00692 | 0.924 | -0.000131 | 0.000524 | 1.00 |

**Table S4:**
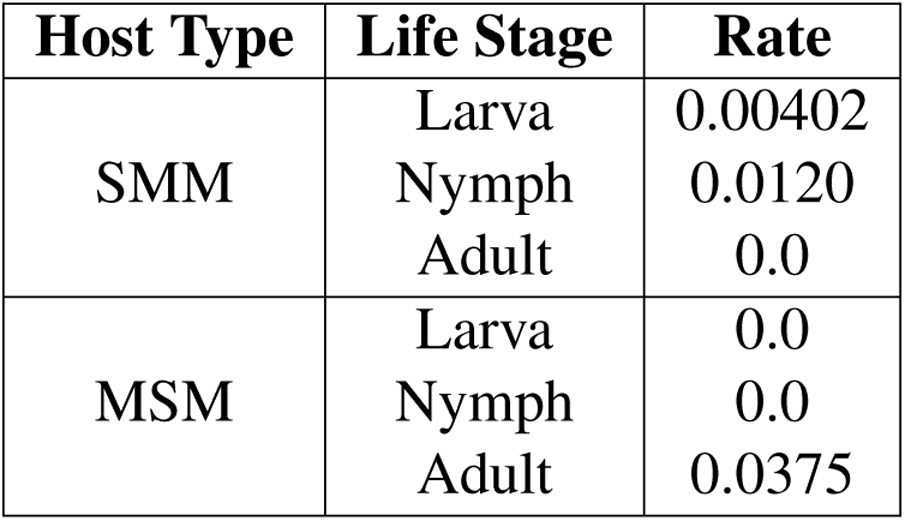
Base Host Finding Rates by Host Type and Life Stage.

**Table S5:**
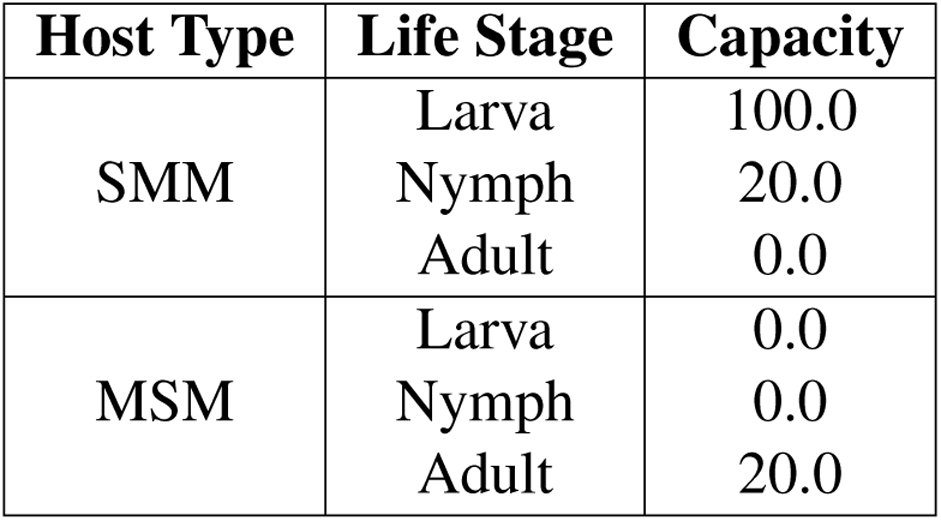
Carrying Capacity of Ticks on Hosts.

| **Host Type** | **Life Stage** | **Capacity** |
|---|---|---|
| SMM | Larva | 100.0 |
|  | Nymph | 20.0 |
|  | Adult | 0.0 |
| MSM | Larva | 0.0 |
|  | Nymph | 0.0 |
|  | Adult | 20.0 |

**Table S6:** Fecundity parameters.

| **Parameter** | **Value** |
|---|---|
| *F*<sub>1</sub> | -25.4181 |
| *F*<sub>2</sub> | 1278.87 |
| *F*<sub>3</sub> | -11588.3 |

**Table S7:** Host-finding rate default coefficients for Immature and Adult temperature and day length dependent equations.

| **Life Stage** | **Parameter** | **Value** |
|---|---|---|
| Immature | *P*<sub>1</sub> | -0.008182 |
|  | *P*<sub>2</sub> | 0.245455 |
|  | *P*<sub>3</sub> | -0.84091 |
|  | *P*<sub>4</sub> | -0.021538 |
|  | *P*<sub>5</sub> | 1.0825 |
|  | *P*<sub>6</sub> | -20.2209 |
|  | *P*<sub>7</sub> | 166.562 |
|  | *P*<sub>8</sub> | -510.835 |
| Adult | *Q*<sub>1</sub> | -0.005051 |
|  | *Q*<sub>2</sub> | 0.232347 |
|  | *Q*<sub>3</sub> | -1.67199 |
|  | *Q*<sub>4</sub> | -0.024278 |
|  | *Q*<sub>5</sub> | 1.23049 |
|  | *Q*<sub>6</sub> | -23.2036 |
|  | *Q*<sub>7</sub> | 193.1549 |
|  | *Q*<sub>8</sub> | -599.3686 |

**Table S8:** Results from Larvae on Host (LOH) sensitivity analysis. The values are partial rank correlation coefficients (PRCC) greater than 0.1 and a p-value less than 0.05. *P*_5_ and *Q*_5_ are parameters in the host finding rate equations for immature and adult ticks. F2 is a parameter in the temperature-dependent fecundity equation.

| **Parameter** | **PRCC** |
|---|---|
| Immature host finding rate *P*<sub>5</sub> | 0.270 |
| Adult host finding rate *Q*<sub>5</sub> | 0.245 |
| Base survival for eggs | 0.152 |
| Critical temp threshold for eggs to larvae | 0.144 |
| Fecundity *F*<sub>2</sub> | 0.107 |

**Table S9:** Results from Nymphs on Host (NOH) sensitivity analysis. The values are partial rank correlation coefficients greater than 0.1 and a p-value less than 0.05. *P*_5_ and *Q*_5_ are parameters in the host finding rate equations for immature and adult ticks.

| **Parameter** | **PRCC** |
|---|---|
| Immature host finding rate *P*<sub>5</sub> | 0.250 |
| Adult host finding rate *Q*<sub>5</sub> | 0.197 |
| Base survival for eggs | 0.146 |
| Critical temp threshold for eggs to larvae | 0.134 |

**Table S10:** Results from adult surplus sensitivity analysis. The values are partial rank correlation coefficients greater than 0.1 and a p-value less than 0.05. *P*_5_ and *Q*_5_ are parameters in the host finding rate equations for immature and adult ticks. *F*_2_ is a parameter in the temperature-dependent fecundity equation.

| **Parameter** | **PRCC** |
|---|---|
| Adult host finding rate *Q*<sub>5</sub> | 0.231 |
| Immature host finding rate *P*<sub>5</sub> | 0.179 |
| Fecundity *F*<sub>2</sub> | 0.128 |
| MSM Base HFR Adult *a* | -0.112 |
| Critical temp threshold for eggs to larvae | 0.105 |

